# Perspectives of pharmacy employees on an inappropriate use of antimicrobials in Kathmandu, Nepal

**DOI:** 10.1101/2022.08.23.503116

**Authors:** Nistha Shrestha, Sulochana Manandhar, Nhukesh Maharjan, Devina Twati, Sabina Dongol, Buddha Basnyat, Stephen Baker, Abhilasha Karkey

## Abstract

**Background:** Unregulated antimicrobial use is common in both hospital and community settings of low- and middle-income countries (LMICs). However, discrete data regarding the use/misuse of antimicrobials at pharmacies in LMICs are limited. This study was conducted to understand the knowledge, attitude, and practice of pharmacy employees on antimicrobial dispensing in Nepal.

**Methods:** We conducted a cross-sectional survey using a structured questionnaire on 801 pharmacy employees working in community and hospital pharmacies located in Lalitpur metropolitan city (LMC) of Kathmandu, Nepal between April 2017 and March 2019.

**Results:** Not all respondents replied to all questions asked. A majority (92%, 738/801) of respondents agreed that the demand for non-prescription antimicrobials was common. Sixty nine percent (437/635) of participants responded that they would ask of prescription before dispensing. Suspected respiratory tract infection was the most common reason demanding for non-prescription antimicrobials, identified by 68% (535/792) of respondents. Azithromycin was the most commonly prescribed and sold antimicrobials, as reported by 46%, (361/787) and 48% (377/779) of participants respectively. A majority (87%; 696/800) of respondents agreed on antimicrobial resistance (AMR) to be a global public health threat; 54% (429/796) perceived antimicrobial misuse to be an important driver of AMR, while only 39% (315/801) replied that judicious dispensing of antimicrobials can help curb AMR.

**Conclusion:** Our study revealed that the unfounded dispensing and use of antimicrobials is prevalent among pharmacies in Kathmandu, Nepal. This overreliance on antimicrobials, notably azithromycin, may escalated the burden of AMR. We identified several drivers of inappropriate antimicrobial dispensing practice in pharmacies, which will aid public health authorities in addressing these issues. Further studies considering the role of other stakeholders, such as physicians, veterinarians, general public, and policy makers are required to obtain a more holistic perspectives on the practices of antimicrobial use so to curb the extant AMR crisis.

## Introduction

The global burden of antimicrobial resistance (AMR) is steadily increasing, while the discovery of new antimicrobials has effectively stopped, creating new challenges in therapeutic management of infectious diseases [1]. The therapeutic management of previously readily treatable diseases, such as typhoid fever, is becoming difficult because of rapid emergence of multi-drug resistant (MDR) variants [2]. It is known that use/misuse of antimicrobials is one of the key factors aiding the accelerated evolution, selection, and spread of antimicrobial resistant bacterial pathogens [1]. The rampant and unregulated use of antimicrobials is common practice in community and hospital settings in low- and middle income countries (LMICs) [1,3]. With a high burden of infectious diseases, coupled with poor sanitary infrastructure and poor public hygiene, the LMICs such as Nepal play a major contributory role in the increasing global burden of AMR [3]. However, discrete data on various aspects of antimicrobial use/misuse is seldom reported from such settings.

In Nepal, local community pharmacies often serve as the first point of contact for healthcare seeking individuals. These facilities not only support the purchasing of prescription antimicrobials, but are also a preferred channel for obtaining a presumptive diagnosis of health issues and accessing non-prescription antimicrobials of their own choice, or as advised by the pharmacies. Therefore, pharmacy employees are arguably the key community stakeholders governing the use/misuse of antimicrobials in communities of LMICs. Understanding perspectives of pharmacy employees on various aspects of antimicrobial dispensing is important to understand the community drivers of AMR in a local setting. Few such studies have been reported from Nepal.

We performed this study to fulfill the existing gaps regarding the knowledge, attitude, and practices of hospital and community pharmacies on various aspects of antimicrobial dispensing and use in Kathmandu, Nepal.

## Materials and methods

According to the data of Department of Drug Administration (DDA) Nepal, as of December 2016, there were 747 registered pharmacies in Lalitpur Metropolitan City (LMC) in Kathmandu, Nepal. During the two year study period between April 2017 and March 2019, the pharmacies in community and hospitals in LMC of Kathmandu, Nepal were spatially mapped based on the data from DDA. The pharmacy employees were approached by study enumerators to ask for their willingness to participate in the study. The willing participants were enrolled and enlisted in the study after providing a written informed consent. The GPS coordinates of the pharmacy were also recorded to aid locating in the subsequent visit. In the later pre-informed visit, the pharmacy participants were asked questions primarily regarding their antimicrobial dispensing practices in an interview format based on the pre-tested structured questionnaire with mostly close-ended questions. The responses were recorded on paper based questionnaire accordingly. Not all the respondents answered to all of the questions. The collected data were electronically recorded in a secure database. The entered electronic data were rechecked and verified. At the end of the study, the data were exported from the database to an excel worksheet for a statistical analysis. The descriptive statistics of qualitative variables were expressed in absolute frequency (N) with percentage (%), while that of quantitative variables was calculated as mean ± standard deviation (SD).

## Results

In this study, the pharmacy employees from a total of 801 pharmacies in LMC, Kathmandu agreed to partake in the study. The privately-owned community pharmacies were the most prevalent (53%; 426/801), followed by private pharmacies inside hospital premises (27%; 218/801). Relatively, the representation of government-owned pharmacies was the least (3.6%; 29/801). A majority (59%; 472/796) of pharmacy employees had an authorized license to dispense antimicrobials, and had pharmacy-specific qualification, while 19% (151/796) of respondents had no specific education, but limited pharmacy-related orientation.

### Antimicrobial dispensing practice and challenges faced

The results on responses of participants on various aspects of antibiotics dispensing are given in Table 1. A high proportion of (92%; 738/801) of respondents agreed that patients often request for antibiotics without any medical prescription (Table 1). When asked about their approach on handling the request for non-prescription antimicrobials, 69% (437/635) of respondent replied that asking for prescription would be the first preference, while 63% (446/708) of the respondents answered that they would dispense non-prescription antimicrobials after some basic interrogation. While dispensing antimicrobials right away seemed to be of the least preference for a majority (81%; 526/652) of respondents. The pharmacy respondents were also asked about the challenges they faced in their profession regarding the dispensing of antimicrobials. In terms of the challenges from patient’s aspects, the self-medication was identified as the major problem for most (58%; 448/768) of the respondents. This was followed by the direct demand for higher-end antimicrobials (35%; 266/758), and patients unwilling to undertake counselling (31%; 234/756). In terms of the treatment-specific challenges faced by the respondents, the poly-pharmacy or, simultaneous use of multiple antimicrobials (33%, 227/697), over prescription of antimicrobials (38%, 268/704), irrational prescribing (35%, 227/657), and frequent use of higher-end antimicrobials (31%, 221/706) were subsequently identified as commonly faced challenges.

**Table 1.**
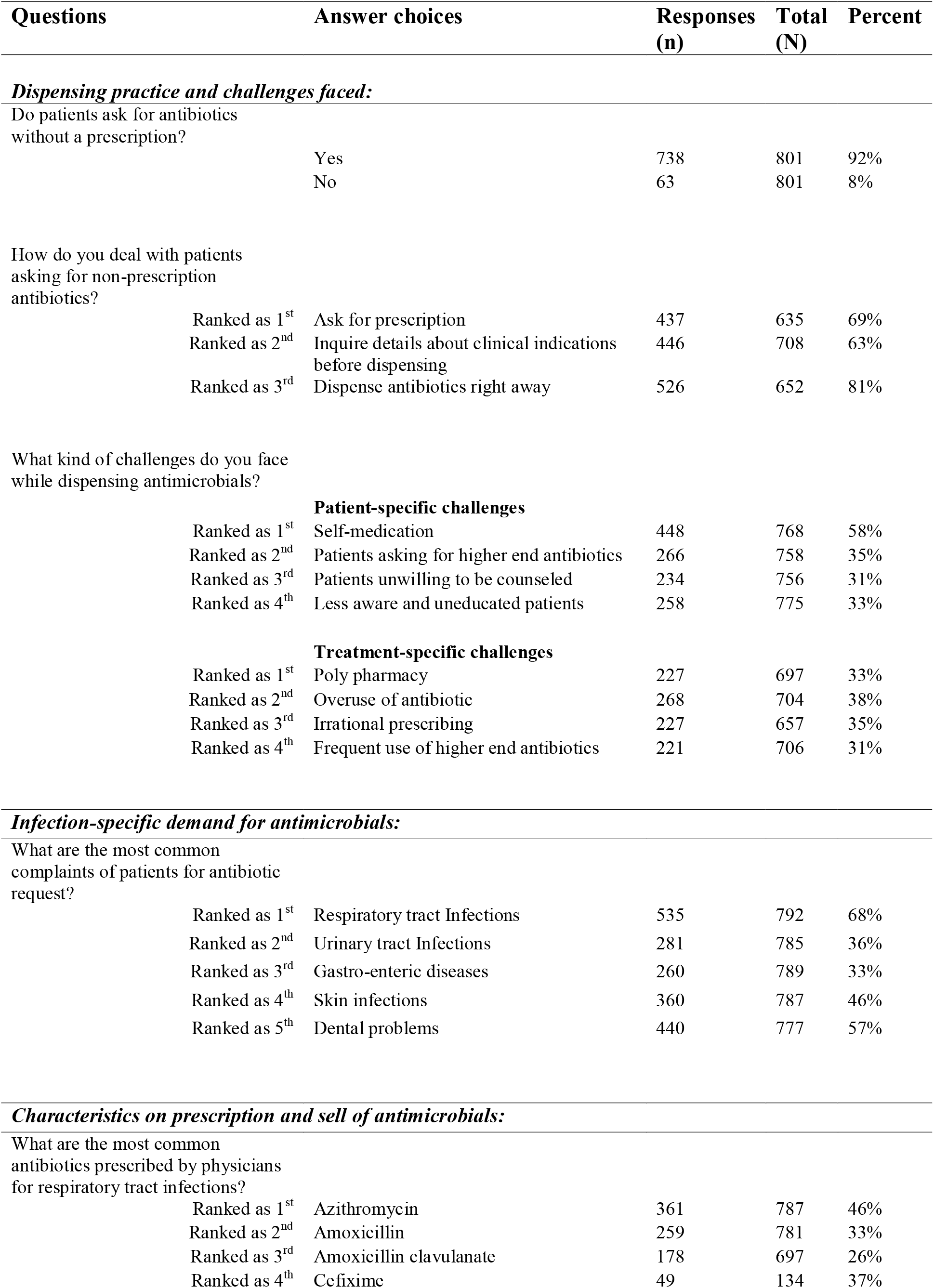
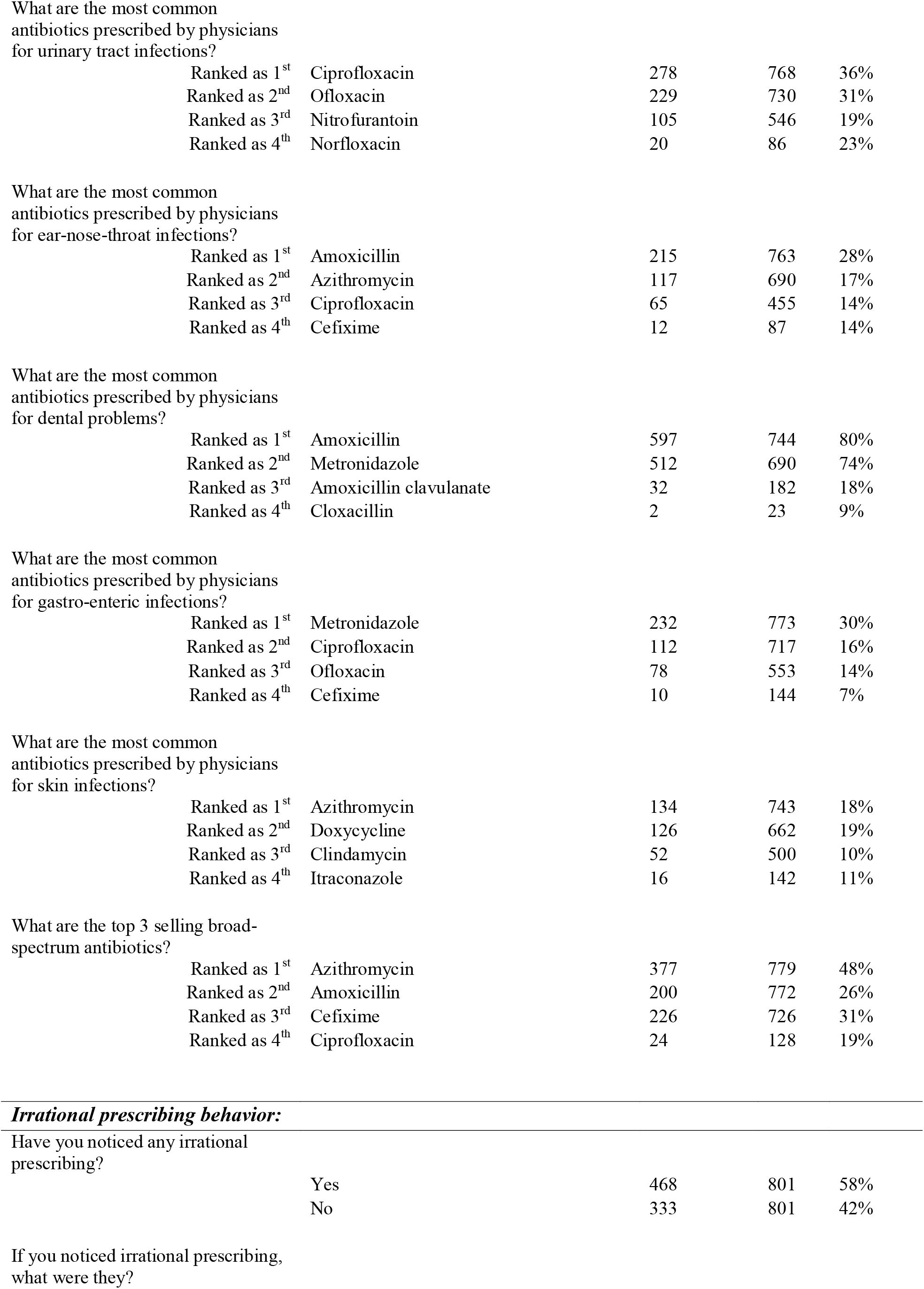
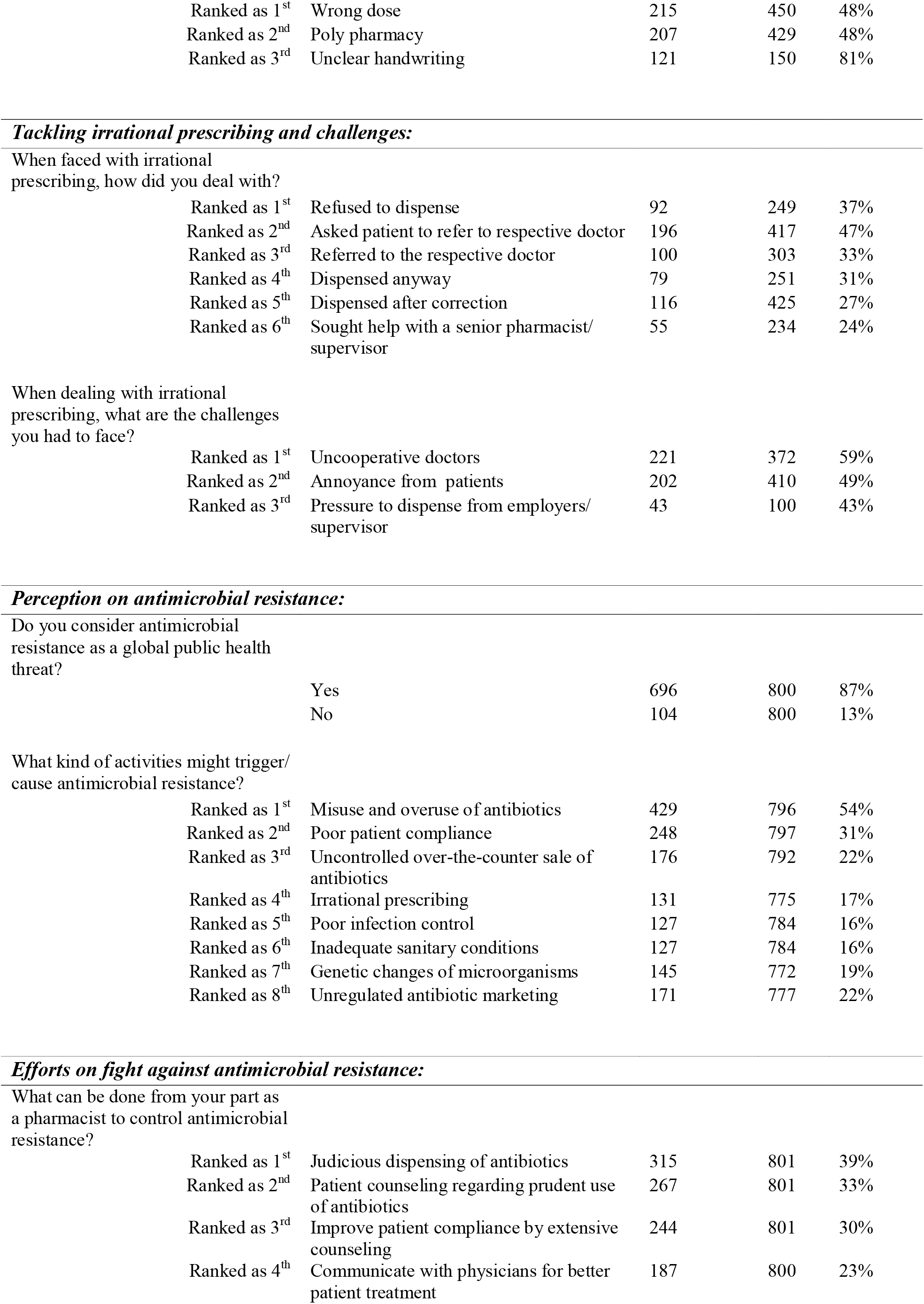
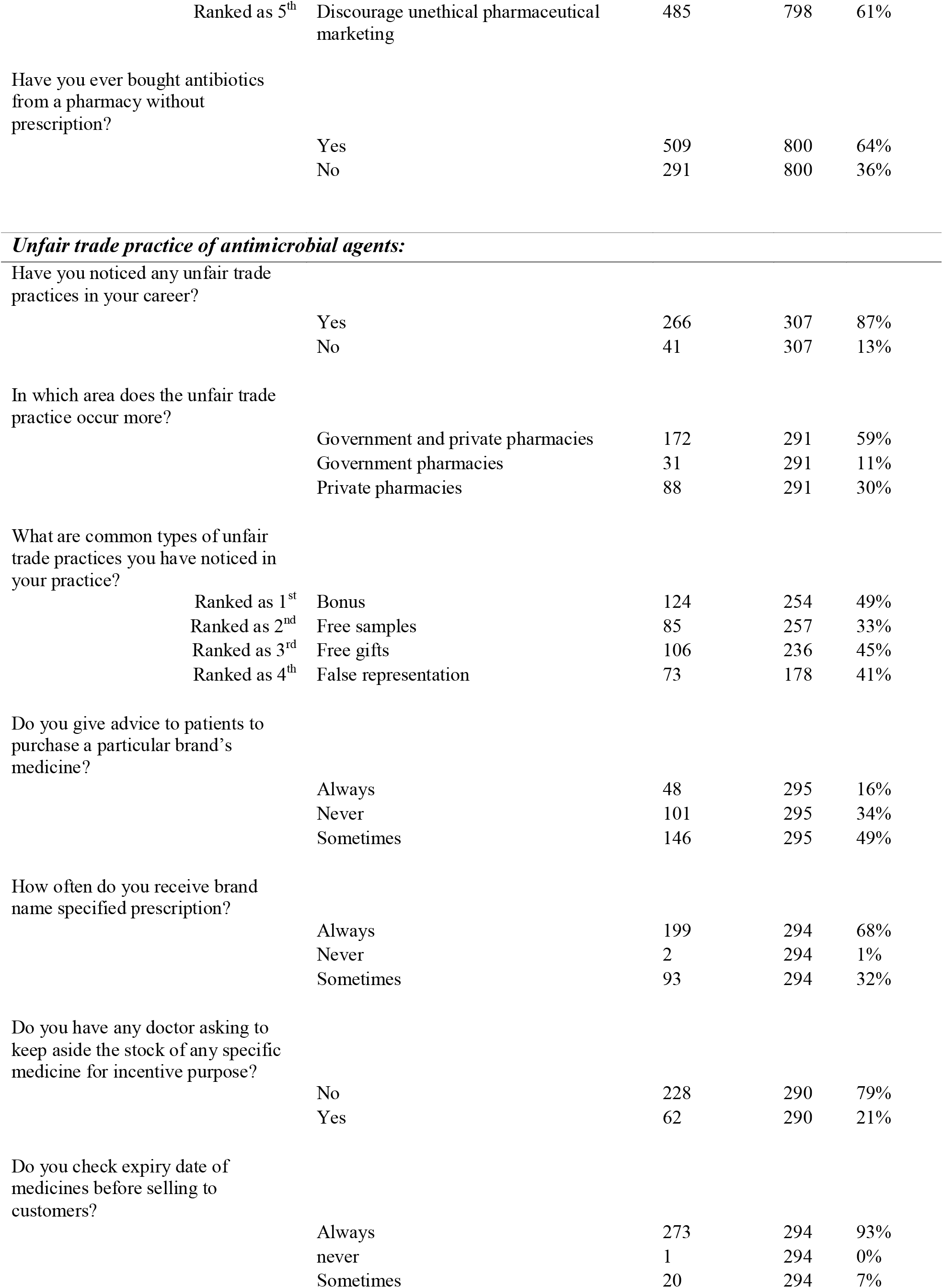
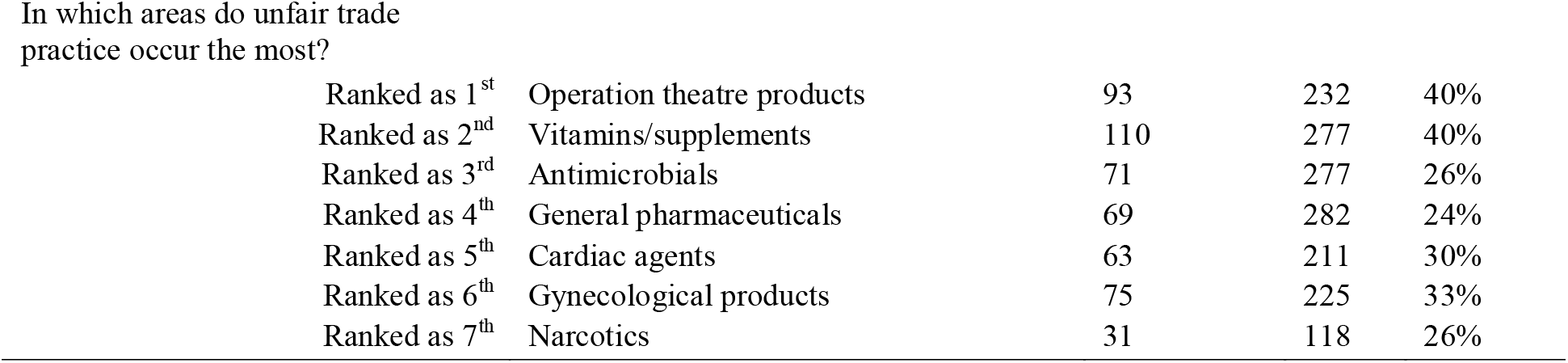
Response of the participating pharmacy employees on various questions related to their knowledge, attitude and practice on antibiotic dispensing

| Questions | Answer choices | Responses (n) | Total (N) | Percent |
| --- | --- | --- | --- | --- |
| ***Dispensing practice and challenges faced:*** |  |  |  |  |
| Do patients ask for antibiotics without a prescription? | Yes | 738 | 801 | 92% |
|  | No | 63 | 801 | 8% |
| How do you deal with patients asking for non-prescription antibiotics? |  |  |  |  |
|  | Ranked as 1<sup>st</sup> Ask for prescription | 437 | 635 | 69% |
|  | Ranked as 2<sup>nd</sup> Inquire details about clinical indications before dispensing | 446 | 708 | 63% |
|  | Ranked as 3<sup>rd</sup> Dispense antibiotics right away | 526 | 652 | 81% |
| What kind of challenges do you face while dispensing antimicrobials? |  |  |  |  |
|  | **Patient-specific challenges** |  |  |  |
|  | Ranked as 1<sup>st</sup> Self-medication | 448 | 768 | 58% |
|  | Ranked as 2<sup>nd</sup> Patients asking for higher end antibiotics | 266 | 758 | 35% |
|  | Ranked as 3<sup>rd</sup> Patients unwilling to be counseled | 234 | 756 | 31% |
|  | Ranked as 4<sup>th</sup> Less aware and uneducated patients | 258 | 775 | 33% |
|  | **Treatment-specific challenges** |  |  |  |
|  | Ranked as 1<sup>st</sup> Poly pharmacy | 227 | 697 | 33% |
|  | Ranked as 2<sup>nd</sup> Overuse of antibiotic | 268 | 704 | 38% |
|  | Ranked as 3<sup>rd</sup> Irrational prescribing | 227 | 657 | 35% |
|  | Ranked as 4<sup>th</sup> Frequent use of higher end antibiotics | 221 | 706 | 31% |
| ***Infection-specific demand for antimicrobials:*** |  |  |  |  |
| What are the most common complaints of patients for antibiotic request? |  |  |  |  |
|  | Ranked as 1<sup>st</sup> Respiratory tract Infections | 535 | 792 | 68% |
|  | Ranked as 2<sup>nd</sup> Urinary tract Infections | 281 | 785 | 36% |
|  | Ranked as 3<sup>rd</sup> Gastro-enteric diseases | 260 | 789 | 33% |
|  | Ranked as 4<sup>th</sup> Skin infections | 360 | 787 | 46% |
|  | Ranked as 5<sup>th</sup> Dental problems | 440 | 777 | 57% |
| ***Characteristics on prescription and sell of antimicrobials:*** |  |  |  |  |
| What are the most common antibiotics prescribed by physicians for respiratory tract infections? |  |  |  |  |
|  | Ranked as 1<sup>st</sup> Azithromycin | 361 | 787 | 46% |
|  | Ranked as 2<sup>nd</sup> Amoxicillin | 259 | 781 | 33% |
|  | Ranked as 3<sup>rd</sup> Amoxicillin clavulanate | 178 | 697 | 26% |
|  | Ranked as 4<sup>th</sup> Cefixime | 49 | 134 | 37% |
| What are the most common antibiotics prescribed by physicians for urinary tract infections? |  |  |  |  |
| Ranked as 1<sup>st</sup> | Ciprofloxacin | 278 | 768 | 36% |
| Ranked as 2<sup>nd</sup> | Ofloxacin | 229 | 730 | 31% |
| Ranked as 3<sup>rd</sup> | Nitrofurantoin | 105 | 546 | 19% |
| Ranked as 4<sup>th</sup> | Norfloxacin | 20 | 86 | 23% |
| What are the most common antibiotics prescribed by physicians for ear-nose-throat infections? |  |  |  |  |
| Ranked as 1<sup>st</sup> | Amoxicillin | 215 | 763 | 28% |
| Ranked as 2<sup>nd</sup> | Azithromycin | 117 | 690 | 17% |
| Ranked as 3<sup>rd</sup> | Ciprofloxacin | 65 | 455 | 14% |
| Ranked as 4<sup>th</sup> | Cefixime | 12 | 87 | 14% |
| What are the most common antibiotics prescribed by physicians for dental problems? |  |  |  |  |
| Ranked as 1<sup>st</sup> | Amoxicillin | 597 | 744 | 80% |
| Ranked as 2<sup>nd</sup> | Metronidazole | 512 | 690 | 74% |
| Ranked as 3<sup>rd</sup> | Amoxicillin clavulanate | 32 | 182 | 18% |
| Ranked as 4<sup>th</sup> | Cloxacillin | 2 | 23 | 9% |
| What are the most common antibiotics prescribed by physicians for gastro-enteric infections? |  |  |  |  |
| Ranked as 1<sup>st</sup> | Metronidazole | 232 | 773 | 30% |
| Ranked as 2<sup>nd</sup> | Ciprofloxacin | 112 | 717 | 16% |
| Ranked as 3<sup>rd</sup> | Ofloxacin | 78 | 553 | 14% |
| Ranked as 4<sup>th</sup> | Cefixime | 10 | 144 | 7% |
| What are the most common antibiotics prescribed by physicians for skin infections? |  |  |  |  |
| Ranked as 1<sup>st</sup> | Azithromycin | 134 | 743 | 18% |
| Ranked as 2<sup>nd</sup> | Doxycycline | 126 | 662 | 19% |
| Ranked as 3<sup>rd</sup> | Clindamycin | 52 | 500 | 10% |
| Ranked as 4<sup>th</sup> | Itraconazole | 16 | 142 | 11% |
| What are the top 3 selling broad-spectrum antibiotics? |  |  |  |  |
| Ranked as 1<sup>st</sup> | Azithromycin | 377 | 779 | 48% |
| Ranked as 2<sup>nd</sup> | Amoxicillin | 200 | 772 | 26% |
| Ranked as 3<sup>rd</sup> | Cefixime | 226 | 726 | 31% |
| Ranked as 4<sup>th</sup> | Ciprofloxacin | 24 | 128 | 19% |

|  |  |  |  |
| --- | --- | --- | --- |
| Yes | 468 | 801 | 58% |
| No | 333 | 801 | 42% |
| If you noticed irrational prescribing, what were they? |  |  |  |  |
| Ranked as 1<sup>st</sup> | Wrong dose | 215 | 450 | 48% |
| Ranked as 2<sup>nd</sup> | Poly pharmacy | 207 | 429 | 48% |
| Ranked as 3<sup>rd</sup> | Unclear handwriting | 121 | 150 | 81% |
| Ranked as 1<sup>st</sup> | Refused to dispense | 92 | 249 | 37% |
| Ranked as 2<sup>nd</sup> | Asked patient to refer to respective doctor | 196 | 417 | 47% |
| Ranked as 3<sup>rd</sup> | Referred to the respective doctor | 100 | 303 | 33% |
| Ranked as 4<sup>th</sup> | Dispensed anyway | 79 | 251 | 31% |
| Ranked as 5<sup>th</sup> | Dispensed after correction | 116 | 425 | 27% |
| Ranked as 6<sup>th</sup> | Sought help with a senior pharmacist/supervisor | 55 | 234 | 24% |
| When dealing with irrational prescribing, what are the challenges you had to face? |  |  |  |  |
| Ranked as 1<sup>st</sup> | Uncooperative doctors | 221 | 372 | 59% |
| Ranked as 2<sup>nd</sup> | Annoyance from patients | 202 | 410 | 49% |
| Ranked as 3<sup>rd</sup> | Pressure to dispense from employers/supervisor | 43 | 100 | 43% |

|  |  |  |  |
| --- | --- | --- | --- |
| Yes | 696 | 800 | 87% |
| No | 104 | 800 | 13% |
| What kind of activities might trigger/cause antimicrobial resistance? |  |  |  |  |
| Ranked as 1<sup>st</sup> | Misuse and overuse of antibiotics | 429 | 796 | 54% |
| Ranked as 2<sup>nd</sup> | Poor patient compliance | 248 | 797 | 31% |
| Ranked as 3<sup>rd</sup> | Uncontrolled over-the-counter sale of antibiotics | 176 | 792 | 22% |
| Ranked as 4<sup>th</sup> | Irrational prescribing | 131 | 775 | 17% |
| Ranked as 5<sup>th</sup> | Poor infection control | 127 | 784 | 16% |
| Ranked as 6<sup>th</sup> | Inadequate sanitary conditions | 127 | 784 | 16% |
| Ranked as 7<sup>th</sup> | Genetic changes of microorganisms | 145 | 772 | 19% |
| Ranked as 8<sup>th</sup> | Unregulated antibiotic marketing | 171 | 777 | 22% |
| Ranked as 1<sup>st</sup> | Judicious dispensing of antibiotics | 315 | 801 | 39% |
| Ranked as 2<sup>nd</sup> | Patient counseling regarding prudent use of antibiotics | 267 | 801 | 33% |
| Ranked as 3<sup>rd</sup> | Improve patient compliance by extensive counseling | 244 | 801 | 30% |
| Ranked as 4<sup>th</sup> | Communicate with physicians for better patient treatment | 187 | 800 | 23% |
| Ranked as 5<sup>th</sup> | Discourage unethical pharmaceutical marketing | 485 | 798 | 61% |
| Have you ever bought antibiotics from a pharmacy without prescription? |  |  |  |  |
|  | Yes | 509 | 800 | 64% |
|  | No | 291 | 800 | 36% |
| ***Unfair trade practice of antimicrobial agents:*** |  |  |  |  |
| Have you noticed any unfair trade practices in your career? |  |  |  |  |
|  | Yes | 266 | 307 | 87% |
|  | No | 41 | 307 | 13% |
| In which area does the unfair trade practice occur more? |  |  |  |  |
|  | Government and private pharmacies | 172 | 291 | 59% |
|  | Government pharmacies | 31 | 291 | 11% |
|  | Private pharmacies | 88 | 291 | 30% |
| What are common types of unfair trade practices you have noticed in your practice? |  |  |  |  |
| Ranked as 1<sup>st</sup> | Bonus | 124 | 254 | 49% |
| Ranked as 2<sup>nd</sup> | Free samples | 85 | 257 | 33% |
| Ranked as 3<sup>rd</sup> | Free gifts | 106 | 236 | 45% |
| Ranked as 4<sup>th</sup> | False representation | 73 | 178 | 41% |
| Do you give advice to patients to purchase a particular brand's medicine? |  |  |  |  |
|  | Always | 48 | 295 | 16% |
|  | Never | 101 | 295 | 34% |
|  | Sometimes | 146 | 295 | 49% |
| How often do you receive brand name specified prescription? |  |  |  |  |
|  | Always | 199 | 294 | 68% |
|  | Never | 2 | 294 | 1% |
|  | Sometimes | 93 | 294 | 32% |
| Do you have any doctor asking to keep aside the stock of any specific medicine for incentive purpose? |  |  |  |  |
|  | No | 228 | 290 | 79% |
|  | Yes | 62 | 290 | 21% |
| Do you check expiry date of medicines before selling to customers? |  |  |  |  |
|  | Always | 273 | 294 | 93% |
|  | never | 1 | 294 | 0% |
|  | Sometimes | 20 | 294 | 7% |
| Ranked as 1<sup>st</sup> | Operation theatre products | 93 | 232 | 40% |
| Ranked as 2<sup>nd</sup> | Vitamins/supplements | 110 | 277 | 40% |
| Ranked as 3<sup>rd</sup> | Antimicrobials | 71 | 277 | 26% |
| Ranked as 4<sup>th</sup> | General pharmaceuticals | 69 | 282 | 24% |
| Ranked as 5<sup>th</sup> | Cardiac agents | 63 | 211 | 30% |
| Ranked as 6<sup>th</sup> | Gynecological products | 75 | 225 | 33% |
| Ranked as 7<sup>th</sup> | Narcotics | 31 | 118 | 26% |

### Infection-specific demand for antimicrobials

The pharmacy respondents were also asked about the most common types of clinical presentations for which they were requested to provide antimicrobials. Sixty eight percent (535/792) of respondents said that the clinical symptoms indicative of respiratory tract infections was the most prevalent complaint associated with a request for antimicrobials, while the clinical signs suggesting urinary tract infections (36%; 281/785), gastrointestinal infections (33%; 260/789), and dental infections (57%; 440/777) were prevalent too.

### Characteristics on prescription and sell of antibiotics

To understand the most frequent types of prescription antimicrobials consumed in Kathmandu Nepal, the respondents were asked questions to identify the most frequent antimicrobials prescribed by physicians by broader infection type. The pharmacy employees ranked azithromycin (46%; 361/787), amoxicillin (33%; 259/781), amoxicillin-clavulanate (26%, 178/697), and cefixime (37%; 49/134) as the four most prescribed antimicrobials for respiratory tract infections. Similarly, for urinary tract infections, over a third of respondents (36%; 278/768) ranked ciprofloxacin to be the first mostly prescribed antimicrobials, followed by ofloxacin (31%; 229/730), and nitrofurantoin (19%; 105/546). Further, the pharmacy respondents were also asked to rank the three most sold broad-spectrum antimicrobials. Azithromycin was ranked as the first most commonly sold antimicrobials identified by almost half (48%, 377/779) of the respondents, followed by amoxicillin (26%, 200/772) and cefixime (31%, 226/726).

### Irrational prescribing behavior

Fifty eight percent (468/801) of respondents replied that they had faced problems of irrational prescribing by medical doctors. Incorrect dosing (47%, 215/450), polypharmacy (48%, 207/431), and illegible writing (80%, 121/150) were ranked to be the first through third most common types of unreasonable prescribing.

### Practices to tackle the irrational prescribing and challenges faced while dealing with it

While 37% (92/249) of respondents answered refusal to dispense to be their first choice of action, 47% (196/417) of respondents replied that requesting patients to obtain a clinical justification before dispensing non-prescription antimicrobials would be their subsequent preference. Similarly, regarding the challenges faced by the pharmacy employees, uncooperative behavior of physicians was identified to be the first most common hurdle faced by 59% (221/372) of the respondents.

### Perception on antimicrobial resistance

A majority (87%; 696/800) of respondents agreed that AMR is a global public health threat; however, 13% (104/800) of respondents did not consider AMR to be a global threat. When asked about their knowledge on the factors contributing to AMR, 54% (429/796) of pharmacy employees identified the misuse and over-use of antimicrobials to be the most important factors driving AMR. Poor patient compliance (31%; 248/797), uncontrolled over-the-counter sale of antimicrobials (22%; 176/792), irrational prescribing (17%; 131/775), poor infection control and inadequate sanitary conditions (16%; 127/784), and the unregulated marketing of antimicrobials (22%; 171/777) were also indicated to be subsequent additional factors escalating AMR.

### Efforts on fighting against antimicrobial resistance

The judicious dispensing of antimicrobials was identified to be the most important factor to curb AMR by many (39%; 315/801) of the respondents. Counselling of the patients about the prudent use of antimicrobials (33%; 267/801), improving antimicrobial compliance among patients (31%; 244/801), and communicating with physicians for better patient treatment (23%; 187/800) were identified as potential solutions in the fight against AMR. Additionally, respondents were also asked about their own personal attitude towards the use of antimicrobials; 64% (509/800) of participants responded that they had bought antimicrobials without prescription for their own personal use.

### Unfair trade practice of antimicrobial agents

When asked if they have ever observed unethical commercial practices in their pharmacy, 87% (266/307) of pharmacy employees agreed on this. Fifty nine percent (171/291) of respondents answered that malpractice was equally prevalent in both government- and privately-owned pharmacies. Forty nine percent (124/254) of respondents suggested that the bonus provided by specific pharmaceutical companies to entice the sale of specific brands of antimicrobials was the most common malpractice they have observed. When respondents were asked if they ever advised clients to buy a specific brand of medicine, half (146/295) of them answered that they often do so. In response to the question regarding the demand for a specific brand of antimicrobials, 68% (199/294) of pharmacy employees replied that they always receive the prescriptions stipulating a specific brand name of antimicrobials; while just <1% (0.7%, 2/294) of respondents denied this activity.

When respondents were asked if they have ever been asked by prescribing clinicians on maintaining a stock of specific brands of medicine that are associated with incentives, a majority (79%, 228/290) of respondents denied that it was the case. Most respondents (93%, 273/294) mentioned that they always check the expiry date of medicines before dispensing them. When the respondents were asked to prioritize the types of pharmaceutical products in which the highest level of unfair trade practice exist, the antimicrobials (26%, 71/277) were ranked to be the third most prevalent pharmaceutical product facing such malpractice, with operation theater items (40%; 93/232), and vitamins/supplements (39%; 110/277) being the first and second ranked items, respectively.

## Discussion

In this prospective cross-sectional survey conducted among 801 pharmacies in LMC of Kathmandu, Nepal, it was evident that inappropriate and irrational prescription of antimicrobial agents is common and widespread in this setting. In high income countries, such as the United States and those in northern Europe, the use of antimicrobials among outpatients is rigorously regulated with strict prescription based dispensing only [4]. However, the use of non-prescription antimicrobials is common in the rest of the world [4]. In fact, in some LMICs, non-prescription antimicrobials are the most commonly sold medicines among prescription-only-drugs [5]. In our study, most (92%, 738/801) pharmacy employees agreed that patients asking for antimicrobials without medical prescriptions was a common situation, which is comparable to a study from Eastern Nepal where 77% of respondents agreed on this [6]. This observation is in contrary to the legislation in Nepal that mandates a strict authorized medical prescription for the dispensing of antimicrobial agents, thereby revealing non-stringent pharmaceutical regulations in Nepal. A systematic review that included 50 studies conducted in 28 developing countries around the world reported that the proportion of antimicrobials dispensed without prescription by pharmacies varied widely, ranging from 3.8% in Tanzania to 100% in Sri Lanka [7].

In LMICs, the role of pharmacies extends beyond dispensing medicines as per the stipulated medical advice. Pharmacies are also engaged in dispensing over-the-counter non-prescription antimicrobials as per patients’ requests. Not just limited to this, the pharmacy operators may even provide their own presumptive diagnosis for sick patients seeking healthcare, and illegally dispense medicines (including antimicrobials) based on their own limited knowledge or experience. Therefore, the pharmacies may play a central role, not only in potentially accelerating the local burden of AMR by directly facilitating inappropriate use of antimicrobials, but also by further jeopardizing the health and safety of general public. Therefore, understanding the knowledge, attitude and practice of pharmacy operators on dispensing antimicrobials is of prime importance. Here, 15% (119/796) of pharmacy operators had neither pharmacy specific education nor training. Previous studies from Nepal (36%, 116/321) and India (75%, 18/24) have also reported on the involvement of unlicensed practitioners in pharmacies, which may directly impact their dispensing practices [6,8]. In our study, 36% of pharmacy respondents agreed that they dispensed non-prescription antimicrobials, which again corresponds with the earlier survey from Nepal [6]. A higher percentage of pharmacy dispensers from India (67%; 174/261) [9] and Pakistan (97%,; 342/353) [10], have also reported to have dispensed non-prescription antimicrobials with or without clarification. In LMICs in South Asia, because of non-stringent government regulations, pharmacies sticking to virtue may instead be at peril of commercial disadvantage. Sixty three percent of pharmacy employees in a study from Nepal believed that refusal to dispense non-prescription antimicrobials would lead to a commercial loss, and nearly 90% agreed that the patients would somehow obtain the non-prescription antimicrobials by attending an alternative pharmacy [6]. Facts such as these might motivate pharmacy employees to deviate from their professional morals. In our study, only half of the respondents perceived that inappropriate use of antimicrobials was the first important factors driving AMR. The earlier study from Nepal reported that nearly 25% of pharmacy respondents declined that inappropriate use of antimicrobials might facilitate AMR [6]. The extent of knowledge, education, and practice as evidenced in this, and other studies from South Asia reflects into the scenario of highly prevalent inappropriate use of antimicrobials in the community.

Acute upper respiratory infections (ARTIs) are considered to be the most common diagnosis for prescription of antimicrobials [11]. As viruses are the most common etiological agents, ARTIs are also associated with an increased burden of inappropriate antimicrobials use and subsequent AMR [11,12]. In our study, 68% of pharmacy respondents replied that respiratory infections were the most common complaints for patients requesting antimicrobials, followed by urinary and gastrointestinal infections. The alternative study from Nepal also reported the highest prevalence of dispensing of antimicrobials were for respiratory (93.3%) and gastrointestinal (91.3%) infections [13]. A study conducted in India reported that 71% (82/115) of pharmacies dispensed antimicrobials without any prescription to the simulated patients presenting with the signs of ARTIs, with 63% (92/115) dispensing antimicrobials for acute gastroenteritis [9]. Further, a study from India reported that pharmacy employees believed that antimicrobials could treat colds (58%, 10/24), viral infections (87%, 20/23), coughs (67%, 16/24), and sore throat (71%, 17/24). Such beliefs and practices among pharmacy dispensers are likely key contributors for inappropriate antimicrobial use in LMICs. In this study, azithromycin was ranked as the first most commonly prescribed antimicrobial for respiratory infections, while ciprofloxacin for urinary infections. Further, azithromycin was also ranked as the first most commonly sold antimicrobials in our study, followed by amoxicillin and cefixime. Koju et al. reported that amoxicillin, azithromycin, amoxi-clavulanate, cefixime, and ciprofloxacin were the top five most commercially promoted antimicrobials for ARTIs in the community pharmacies of Nepal, which could have a direct influence on an increased prescription and sell of these antimicrobials [12]. Another study from Nepal found that cefixime (38%), amoxicillin (29.3%), ciprofloxacin (13.7%), and azithromycin (8.1%) were the most commonly dispensed antimicrobials by community pharmacies [13]. Studies from India and Pakistan have also reported that azithromycin, ciprofloxacin, and cefixime were the most commonly prescribed or sold antimicrobials [9,10]. Azithromycin, cefixime, and ciprofloxacin belong to the watch group antimicrobials under the WHO AWaRe category of antimicrobials, that have increased risk of rendering resistance or toxicity [14]. The finding that these crucial antimicrobials were the most commercially promoted, prescribed, and sold in LMICs portrays a challenging scenario facilitating an escalated emergence and spread of antimicrobial resistance.

The burden of faeco-orally transmitted infectious diseases, such as typhoid, are reflective of sub-standard public health measures, and are highly prevalent in LMICs including Nepal [15]. The emergence, selection, and spread of antimicrobial resistant variants of *Salmonella enterica* serovars Typhi and Paratyphi A have been observed, leading to the subsequent treatment failure [1,2,16–18]. Once easily treatable with first-line antimicrobials such as amoxicillin and cotrimoxazole, the therapeutic management of typhoid is now more challenging, requiring second- and third-line antimicrobials such as azithromycin and parenteral ceftriaxone [16]. A study from Pakistan reported the emergence and spread of extensively drug resistant (XDR) *S*. Typhi harboring genes conferring AMR to all first and second-line antimicrobials along with ceftriaxone, leaving azithromycin as the only widely available oral antimicrobial for treating typhoid [17]. A recent study from Nepal reported clinical isolates of *S*. Typhi that were resistant to azithromycin and ciprofloxacin [18]. We surmise that rampant consumption of azithromycin, cefixime, and ciprofloxacin, as was inferred by their highest prescription and sale in this study, further corroborates the potential scenario of untreatable infectious diseases such as typhoid [16].

## Conclusions

Here, we assessed knowledge, attitude, and practices of pharmacy employees in LMC of Kathmandu, Nepal on various aspects of inappropriate use of antimicrobials. Our study revealed that the demand and without-prescription use of several antimicrobials, including the WHO watch group antimicrobials, is highly prevalent in Kathmandu, Nepal. This scenario may escalate emergence and spread AMR, thereby threatening the therapeutic management of infectious diseases including the otherwise easily treatable ones as evidenced in typhoid fever. Several factors driving inappropriate practice among pharmacy employees were also identified which may aid in formulating and strengthening policies at a local level. Further studies are required to understand the perspectives and practices of other functional stakeholders of the community such as prescribing physicians, veterinarians, medicine dealers, policy makes, other concerned authorities, and general population. Results from such will help gather multi-sectorial information on antimicrobial demand, prescription, sale and consumption to develop holistic strategies to fight against the extant AMR crisis.

## Acknowledgements

We would like to acknowledge all pharmacies and pharmacy employees for participating and providing valuable information on their professional experience and practice. We like acknowledge all enumerators involved in this study for their work on collecting and verifying information from the field.

